# Prediction of white matter hyperintensities evolution one-year post-stroke from a single-point brain MRI and stroke lesions information

**DOI:** 10.1101/2022.12.14.520239

**Authors:** Muhammad Febrian Rachmadi, Maria del C. Valdés-Hernández, Stephen Makin, Joanna Wardlaw, Henrik Skibbe

## Abstract

Predicting the evolution of white matter hyperintensities (WMH), a common feature in brain magnetic resonance imaging (MRI) scans of older adults (i.e., whether WMH will grow, remain stable, or shrink with time) is important for personalised therapeutic interventions. However, this task is difficult mainly due to the myriad of vascular risk factors and comorbidities that influence it, and the low specificity and sensitivity of the image intensities and textures alone for predicting WMH evolution. Given the predominantly vascular nature of WMH, in this study, we evaluate the impact of incorporating stroke lesion information to a probabilistic deep learning model to predict the evolution of WMH 1-year after the baseline image acquisition, taken soon after a mild stroke event, using T2-FLAIR brain MRI. The Probabilistic U-Net was chosen for this study due to its capability of simulating and quantifying the uncertainties involved in the prediction of WMH evolution. We propose to use an additional loss called volume loss to train our model, and incorporate stroke lesions information, an influential factor in WMH evolution. Our experiments showed that jointly segmenting the disease evolution map (DEM) of WMH and stroke lesions, improved the accuracy of the DEM representing WMH evolution. The combination of introducing the volume loss and joint segmentation of DEM of WMH and stroke lesions outperformed other model configurations with mean volumetric absolute error of 0.0092 *ml* (down from 1.7739 *ml*) and 0.47% improvement on average Dice similarity coefficient in shrinking, growing and stable WMH.

## Introduction

### White matter hyperintensities and their progression

White matter hyperintensities (WMH) are one of the main neuroradiological features of cerebral small vessel disease (SVD) and have been commonly associated with stroke, aging, and dementia progression^1–3^. They are often observed in T2-weighted and T2-fluid attenuated inversion recovery (T2-FLAIR) brain magnetic resonance images (MRI), appearing as bright regions. Small subcortical infarcts may be indistinguishable from WMH on structural MRI in absence of intravenous contrast due to sharing similar image intensity characteristics^4^, and if mistaken for WMH could negatively impact the design of clinical research trials^5^.

Clinical studies have indicated that some patients exhibit WMH progression over time (i.e., increasing in volume)^6–8^ while some show WMH regression over time (i.e., shrinking in volume)^9,10^, although these are fewer in proportion compared to those reporting an increase in volume^10^. Another study indicated that WMH dynamically change over time with clusters of WMH individually shrinking, staying unchanged (i.e., stable), or growing, these being observed at the same time point within the same individual^11^. These variations have been associated with patients’ comorbidities and clinical outcome^3,12^. A meta-analysis on rate and risk factors for WMH volume growth specifically, concluded that these vary with the characteristics of the sample, although hypertension, age, baseline WMH volume and smoking seemed to be the main contributors^13^. And a growing number of clinical studies have indicated that, in addition to age^8^, previous strokes^14^ and genetics^15–18^ also influence the rate and direction of WMH evolution. But, as one clinical study and another meta-analysis acknowledged, current knowledge about factors influencing WMH evolution is still incomplete and poorly understood^3,10^.

Interestingly, despite increasing evidence on WMH burden at baseline being the determinant factor on the rate and magnitude of WMH progression (and regression)^13^, increase in WMH volume has been found to be a better predictor of persistent cognitive impairment (i.e., a potential precursor to Alzheimer or vascular dementia) than baseline WMH burden^19^. However, evidence that overall reduction of WMH volume over time can prevent functional decline is scarce^2^. In terms of spatial WMH evolution, a study on patients that had a mild stroke of type lacunar found that post-stroke cognition at 1 and 3 years was affected by the location of WMH^20^. But despite evidence on the importance and benefit of studying WMH spatial distribution^21^, there are limited approaches to predict spatial WMH evolution. Predicting the evolution of WMH is crucial for understanding the dynamics of small vessel disease and ultimately provide better care and prognosis for individual patients, but it remains a difficult task because of the different rate and direction of the evolution of individual WMH clusters and their interplay with other imaging features of vascular disease and brain parenchymal changes^14^. Specifically, 1 year after stroke, reported WMH changes are mild^13^, thus posing an additional challenge for their accurate identification.

### Precedent work in estimating WMH evolution

Despite the high accuracy displayed by several fully-automatic deep learning schemes segmenting WMH^22^, most of the algorithms applied in longitudinal studies on WMH evolution have been so far semi-automatic^13^. Various deep learning models have been proposed to predict the spatial evolution of WMH^23–25^. These studies, have represented WMH spatial evolution by a map called *disease evolution map* (DEM) which indicates the WMH voxels that shrink, grow, or remain stable at a further time point. DEM can be generated by subtracting images of manually labeled WMH from different time points. Previous studies generated the DEM by subtracting a baseline image of semi- or fully-automatically labeled WMH of a patient (Visit 1, V1) from a follow-up image of semi- or fully-automatically labeled WMH from the same patient one year after (Visit 2, V2)^24,25^. An example of DEM is visualised in Figure 1B.

**Figure 1.**
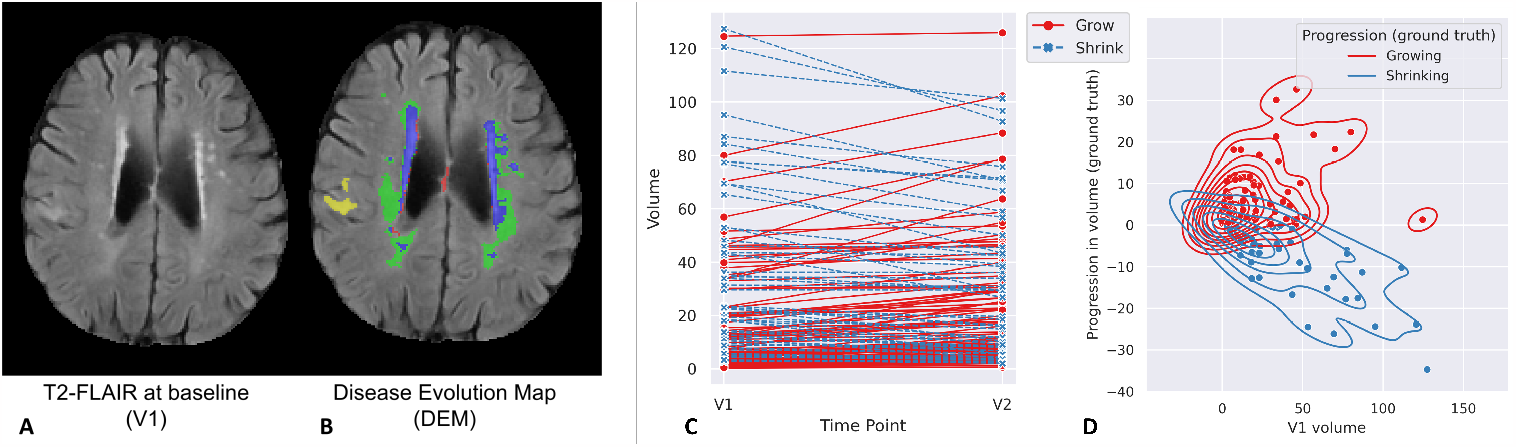
**A:** Brain-extracted FLAIR axial slice of the baseline scan or V1. **B:** Visualisation of disease evolution map (DEM) of white matter hyperintesities (WMH). Red represents shrinking WMH, green represents growing WMH, blue represents stable WMH, and yellow represents stroke lesions. **C:** Volumetric progression of WMH (in *ml*) from V1 to V2 (1 year apart) for all subjects from our dataset. **D:** shows the distribution of volumetric progression of WMH (in *ml*) based on WMH volume at V1 for all subjects.

A recently proposed model for predicting the DEM of WMH based on a Probabilistic U-Net^26^, generates multiple DEM predictions for a single brain MRI data^25^. This model was proposed to solve the challenge of representing spatial uncertainty^24^, given difficulties in distinguishing intensities and textures of shrinking and growing WMH in T2-FLAIR brain MRI. Models using Probabilistic U-Net performed significantly better than the classical U-Net models in predicting the evolution of WMH using DEM^25^.

All these previous approaches have focused, almost exclusively, on the image modality as input and the appearance of WMH themselves, ignoring other clinically relevant factors. A subsequent study incorporated volume of stroke lesions as auxiliary input to the prediction model, but it did not improve the prediction results^24^. Another study^27^ used radiomic signatures of the normal-appearing tissue as auxiliary variables to vascular risk factors in a logistic regression model to predict general “progression” vs. “no progression” of WMH, and reported that radiomics improved the accuracy of the model by approximately 10%, but did not analyse the spatial change of WMH. Thus, incorporating clinically associated factors into the predictive model remains a challenge for estimating the spatial evolution of WMH.

### Related Approaches

Studies that develop predictive models for disease progression from medical image modalities using machine/deep learning can be categorised, generally, into the three different approaches listed below.

1. **Approaches predicting the outcomes of a disease**. These approaches are commonly used for diseases with high rates of mortality and disability. Some examples are those predicting the outcomes of COVID-19^28^, multiple sclerosis^29^, and traumatic brain injury^30,31^.
2. **Approaches predicting the progression of a disease with regards to the pathological timeline and/or commonly associated disease markers**. These approaches are commonly used for diseases with multiple stages of development and which take time to progress, such as dementia and Alzheimer’s Disease (AD), with mild cognitive impairment (MCI) being their transitional stage^32^. Some examples are predicting conversion of MCI patients to AD^33^, conversion of healthy individuals to MCI and AD^34^, and predicting the progression of multimodal AD markers (e.g., ventricular volume, cognitive scores, etc.)^35^.
3. **Approaches predicting dynamic changes (evolution) of specific disease features**. These approaches model and predict spatial changes of specific disease features such as evolution of WMH, enlargement of ventricles, and brain atrophy. Other examples are predicting lung nodule progression of pulmonary tumour^36^, predicting dynamic change of brain structures from healthy individuals to MCI and AD patients^37^, and studies for predicting the evolution of WMH in brain images of stroke patients^23–25^

This study belongs to the third category, in which a predictive model is used to spatially estimate the dynamic changes of WMH on an MRI scan at a certain time point. This third category is the most challenging because of the complexity and resolution of the data/image being predicted, especially when the time-point estimated is close to the baseline scan. While approaches in the first and second categories predict classes which are the disease outcomes (e.g., survive, death), classes of disease stages (e.g., MCI, AD), or associated disease markers (e.g., age, cognitive scores) from medical imaging data, approaches in the third category predict the evolution of disease’s imaging features (e.g., lesions and their volumes) spatially, i.e., throughout the entire image space.

### Our Contributions

The **main contributions** of this study are two-fold, and show that they considerably improve the prediction of WMH volume and spatial change 1 year after a mild-to-moderate stroke event:

1. **incorporating stroke lesions’ information to the prediction model** and
2. **adding a volume loss to the cost function** (formulated as the mean squared error between the predicted and the reference future WMH volumes) to improve prediction of WMH evolution voxel-wise.

As part of a comprehensive set of evaluations, We also evaluate the output from our schemes against the clinical visual scores for WMH evolution^38^, and analyse the degree of uncertainty in our predictions.

## Proposed Deep Learning Model

Uncertainties are unavoidable when predicting the progression of WMH. Previous studies showed that incorporating uncertainties into a deep learning model, either by incorporating Gaussian noise as auxiliary input^24^ or using a conditional variational autoencoder in the shape of a Probabilistic U-Net with adversarial training^25^, improved prediction results, thus justifying the use of a Probabilistic U-Net with adversarial training in the present study.

### Probabilistic U-Net with adversarial training

The uncertainty associated with the randomness in the dynamism of the WMH clusters is commonly known as *aleatoric uncertainty*^41^. It constitutes the biggest challenge in predicting WMH evolution, due to differences between experts in WMH delineation (i.e., ground truth reliability issues), and difficulty in differentiating textures and intensities of shrinking and growing WMH in the T2-FLAIR MRI sequence^24^. This uncertainty cannot be reduced by simply adding more training data^41^. The use of a Bayesian deep learning model named Probabilistic U-Net^26^ was previously proposed to overcome this challenge, and generated better prediction results than non-probabilistic models^25^. In this study, we modify the previously proposed approach, as Figure 2A schematically illustrates.

**Figure 2.**
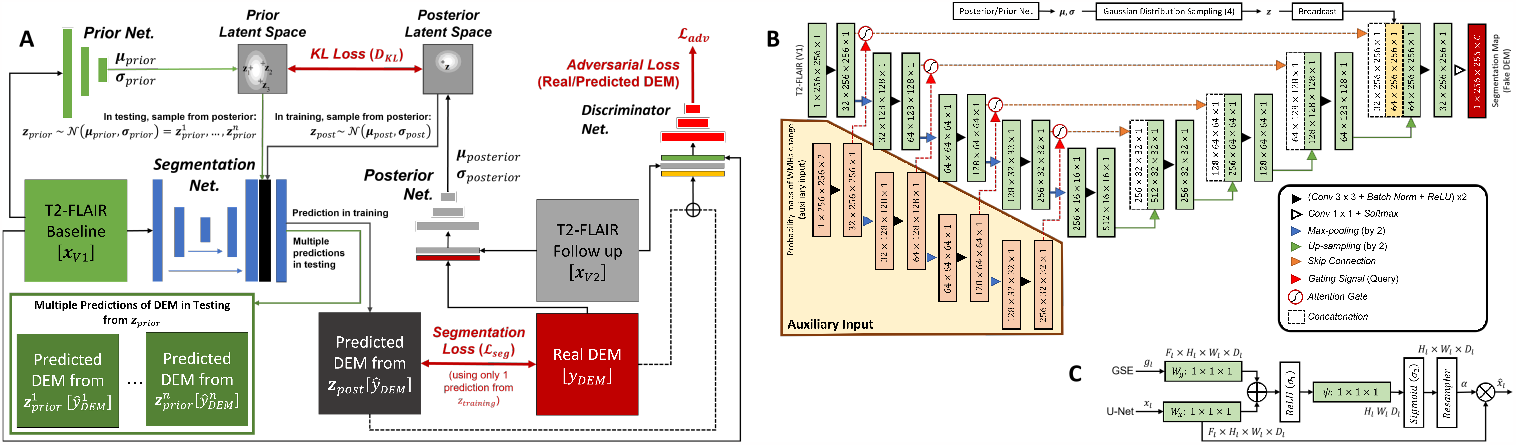
**A:** Schematic representation of the Probabilistic U-Net^26^ with adversarial training^39^ used in this study, firstly introduced in a previous work^25^. **B:** Segmentation network of Probabilistic U-Net used in this study, which is based on the original U-Net extended into Attention U-Net only when probability maps of WMH change are used as auxiliary input. The output channel of *C* is either 5 or 4 depending on whether stroke lesions are jointly segmented or not, respectively. **C:** Schematic of additive attention gate (AG) used in this study, firstly introduced in^40^. Input features (*x*_*l*_) are from the U-Net’s skip connection while gating signals (*g*_*l*_) are from the gating signal encoder (GSE). Attention coefficients (*α*) are learned in the training process and used to scale input features *x*_*l*_ to highlight important areas.

The probabilistic U-Net with adversarial training consists of a U-Net configuration^42^, two variational encoders called Prior Net and Posterior Net, and a discriminator network for adversarial training. In this study, the U-Net was used as segmentation network for predicting the DEM. Meanwhile, Prior Net and Posterior Net were used for variational inference so that uncertainty in predicting future WMH evolution is modeled probabilistically. Prior Net estimates a low-dimensional Gaussian distribution called prior latent space by producing its mean(s) and variance(s) from T2-FLAIR MRI at baseline (i.e., V1, denoted *x*_*V*1_). Whereas, Posterior Net estimates another low-dimensional Gaussian distribution called posterior latent space by producing its mean(s) and variance(s) from the follow-up T2-FLAIR MRI (i.e., V2, denoted *x*_*V*2_) and ground truth DEM (*y*_*DEM*_). In reality, the posterior latent space is unknown because the follow-up T2-FLAIR MRI and the ground truth DEM are not present. Because of that, *Kullback-Leibler* divergence is used during training to estimate a posterior latent space from the prior latent space, which is obtained from the baseline T2-FLAIR MRI. In training, a sample ***z***_*post*_ is taken from the posterior latent space (***z***_*post*_ *∼ 𝒩*(***μ*** _*post*_, ***σ***_*post*_) and then broadcasted and concatenated to the segmentation network. Multiple predictions of _*DEM*_ 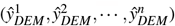 can be generated by using multiple samples 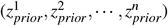 from the prior latent space (***z***_*prior*_ *∼ 𝒩*(***μ*** _*prior*_, ***σ***_*prior*_). In this study 30 different DEM (_*DEM*_) predictions were generated from 30 samples of *z*_*prior*_ from Prior Net for each input data/patient in the inference, and then averaged to get the final DEM prediction. Lastly, a discriminator network is used for adversarial training to enforce anatomically realistic DEM with regards to the T2-FLAIR MRI at V1 and V2, similar to previous work^24^. The discriminator network has 13.3M trainable parameters.

### Incorporation of stroke lesions information

Clinical studies have indicated that there are strong correlations between stroke occurrence and progression of WMH over time^14^. In a previous study^24^, stroke lesion volume was used as an auxiliary input to a framework designed to estimate WMH evolution, but it was outperformed by the use of Gaussian noises as auxiliary input representing uncertainty. Thus, in this study, we explore how information on stroke lesions can be incorporated to the probabilistic framework for better prediction of WMH evolution. We propose two different approaches: 1) jointly segmenting the WMH DEM and stroke lesions, and 2) incorporating probabilistic maps of WMH change in relation to stroke lesions’ locations. The second proposed approach is more complex than the first proposed approach because it needs multiple preprocessing steps.

### Joint segmentation of DEM and stroke lesions

Due to the similar tissue signal intensity of WMH and ischaemic stroke lesions in T2-FLAIR brain MRI, we hypothesised that performing a joint segmentation of the WMH DEM and stroke lesions will improve the accuracy in the prediction of the WMH DEM because the deep learning model will automatically learn the spatial correlation between both features. In this proposed approach stroke lesions do not need to be excluded in the preprocessing steps like in preceding works^24,25^. This approach can be implemented by adding an output channel to the segmentation layer of the segmentation network, thus increasing the number of output channels from the originally four (i.e., channels for background, shrinking WMH, growing WMH, and stable WMH), to five channels. Note in Figure 1B that the label ‘stroke lesions’ has been added to the DEM of WMH. In this setting, the generator has 31.5M trainable parameters.

### Probabilistic maps of WMH change in relation to stroke lesions’ locations

Results from a clinical study indicate that there are strong correlations between stroke lesions’ location at baseline (V1) and WMH evolution after 1 year (V2)^20^ for patients with a stroke of type lacunar. Specifically, if stroke lesions are subcortical and located in either the *centrum semiovale* or the *lentiform nucleus* at V1, then there are significant changes to the WMH at V2 (both in volume and location) specific to the location of the stroke lesions at V1. This clinical study made available probability maps of WMH change indicating brain locations where changes of WMH are significant at V2 depending on the infarcted region after accounting for vascular risk factors (VRF)^43^.

We use these probability maps as auxiliary data input to an attention U-Net^40^ within the Probabilistic U-Net’s segmentation network. In it, the information of the probability maps is encoded through the gating signal encoder (GSE), with outputs used as gating signal in multiple resolutions (see Figure 2B). The general idea of this approach is to focus the attention of the segmentation network on the areas that have high probability of WMH change according to the locations of the stroke lesions. For this approach, we performed brain parcellation and registration of the probability maps (in standard image space) to each patient’s space to identify the locations of stroke lesions for each specific patient. In this setting, the generator has 32.2M trainable parameters.

Similar to the original Attention U-Net^40^, this study uses an additive attention gate (AG), but obtains the gating signals from the GSE instead of from the outputs of the next (coarser) convolutional block. The schematic of the additive AG can be seen in Figure 2C. Input features (*x*_*l*_) are from the U-Net’s skip connections, gating signals (*g*_*l*_) are from the gating signal encoder (GSE), *α* are the attention coefficients learned in the training process used to scale input features *x*_*l*_ to highlight important areas, ⊕ is an element-wise addition, ⊗ is an element-wise multiplication, and *W*_*g*_, *W*_*x*_, and *Ψ* are 1 *×* 1 *×* 1 convolution operations.

### Configurations of the proposed approach

In this study we evaluate four configurations of the segmentation network. Three different configurations incorporating probabilistic maps of WMH and/or stroke lesions were compared with the vanilla U-Net. All of them took 5-8 minutes to train per epoch.

1. **PUNet**: The original U-Net^42^ is used for the segmentation network.
2. **PUNet-wSL**: The original U-Net^42^ is used for the segmentation network, and joint segmentation of DEM of WMH and stroke lesions is performed.
3. **Att-PUNet**: Attention U-Net with probabilistic maps of WMH change is used for the segmentation network, instead of the original U-Net.
4. **Att-PUNet-wSL**: Attention U-Net with probabilistic maps of WMH change is used for the segmentation network, and, simultaneously, joint segmentation of DEM and stroke lesions is performed.

## Experimental Setting

This section describes the dataset, training scheme, cost function, and evaluation metrics this study uses.

### Dataset

For comparability of our results with those previously published, we use the same dataset as^24^, which comprises MRI data from *n* = 152 patients that had a mild-to-moderate stroke and consented to participate in a study of stroke mechanisms^3^. The study protocols were approved by the Lothian Ethics of Medical Research Committee (REC 09/81101/54) and NHS Lothian R+D Office (2009/W/NEU/14), on the 29th of October 2009. All patients were imaged with the same acquisition protocol at two time points (i.e., baseline scan (V1), and a year after the baseline scan (V2)). In total, 304 MRI from 152 stroke patients (i.e., 152 V1 MRI and 152 V2 MRI) were used. Overall increase in WMH volume was identified in 98 of the 152 patients and reduction of WMH total volume in 54 patients. The magnitudes of WMH change (in *ml*) and their distribution for all patients can be seen in Figure 1C and 1D.

All T2-FLAIR brain MRI were acquired with a GE 1.5T scanner, and a semi-automatic multi-spectral method was used to produce several brain masks including intracranial volume, cerebrospinal fluid, stroke lesions, and WMH, all which were visually checked and manually edited by an expert^44^. For the prediction of WMH evolution from V1 to V2, T2-FLAIR brain MRI at follow-up (V2) and T2-FLAIR brain MRI at baseline (V1) were linearly and rigidly aligned to a common space using FSL-FLIRT^45^. The space transformations were applied to all labels (i.e., binary/indexed masks) including manually-derived (i.e., after manually correcting results from a semi-automatic segmentation) labels of WMH. The spatial resolution of the images was 256 *×* 256 *×* 42 with slice thickness of 0.9375 *×* 0.9375 *×* 4 mm. We generated a DEM for each patient by subtracting the manually corrected segmentation of WMH at V1 from the manually corrected segmentation of WMH at V2.

### Data pre-processing for incorporation of probabilistic maps of WMH change

Given the influence of stroke lesion location in WMH change and evolution patterns when the stroke lesions are located at the *centrum semiovale* or the *lentiform nucleus*^20^, we only used probability maps of WMH change based on stroke lesions incident at *centrum semiovale* or *lentiform nucleus*, publicly available^1^.

Probability maps in the standard space were obtained from a clinical study^20^ and then registered to each patient’s native space using niftyreg through TractoR^46^. To identify the location of stroke lesions within a human brain, an age-relevant brain template and its corresponding brain parcellation), also publicly available^47^, were registered to each patient’s native space. If there were no stroke lesions at *centrum semiovale* or *lentiform nucleus* in a patient, then zero matrices were used as probabilistic maps (i.e., there are no specific areas of the brain or feature maps that the neural networks should look for via attention). Both probabilistic maps for *centrum semiovale* or *lentiform nucleus* were concatenated before being used as auxiliary input in the segmentation network (see Figure 2B for illustration).

### Training scheme

To facilitate comparability between methods and results, we used the same preprocessing pipeline as previous studies^24,25^. To make sure all patients are used in both training and testing, 4-fold cross validation with 512 epochs was performed with each fold consisting of 114 MRI for training and 38 for testing. In the training phase, we randomly chose 14 out of 114 MRI training data for validation and used that for selecting the best model that produced the lowest validation loss (i.e., error difference during training). Values of T2-FLAIR brain MRI were normalised into zero mean and unit variance for each patient. Data augmentations of shifting, scaling, horizontal and vertical flip, and elastic transformations were performed.

### Cost function

We used three lost functions in training to optimize the different networks. These were: 1) segmentation loss (*ℒ*_*seg*_), 2) probabilistic loss using *Kullback-Leibler* Divergence (*D*_*KL*_), and 3) adversarial loss (*L*_*adv*_). We used the segmentation loss to compare the output of the segmentation network (i.e., the predicted DEM segmentation) against the ground truth of the DEM. The probabilistic loss was used to compare the similarity between prior and posterior latent spaces. And the adversarial loss was used to compare the similarity between the ground truth DEM and the predicted DEM.

### Segmentation loss

For the segmentation loss, we used the weighted focal loss with *γ* (i.e., focal loss’ hyperparameter) set to *γ* = 2 following the recommendation of the original paper^48^. Equation 1 describes the weighted focal loss function for all pixels from an MRI slice where *y*_*i,c*_ *∈ {*0, 1*}* indicates the class membership for pixel *i* to class *c, p*_*i*_ the predicted probability that pixel *i* belongs to class *c*, and *α*_*c*_ is the weight for class *c*. The larger the value of *α*_*c*_, the larger the contribution of class *c* to the loss value. *P* is the random variable for the predicted probability, *Y* is the random variable for the target classes, *α* are the weights for all classes, *N* is the number of pixels in an axial MRI slice (i.e., *N* = 256), and *M* is the number of classes in the DEM (i.e., *N* = 4 if stroke lesions are not automatically segmented and *N* = 5 if otherwise). Based on our preliminary experiments, the best weights were *α*_*c*=0_ = 0.25 for background, *α*_*c*=1_ = 0.75 for shrinking WMH, *α*_*c*=2_ = 0.75 for growing WMH, *α*_*c*=3_ = 0.5 for stable WMH, and *α*_*c*=4_ = 0.75 for stroke lesions.

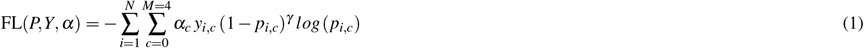

Note that the predicted segmentation of the DEM produced by the Probabilistic U-Net is conditioned to either the posterior or the prior latent space. In training, the predicted DEM segmentation is conditioned to the posterior latent space defined by ***z***_*post*_ *∼ 𝒩*(***μ*** _*post*_, ***σ***_*post*_) and modelled by the Posterior Net. On the other hand, the predicted DEM segmentation is conditioned by the prior latent space that is formulated as ***z***_*prior*_ *∼ 𝒩*(***μ*** _*prior*_, ***σ***_*prior*_) and modelled by the Prior Net in testing/inference. Thus, the probabilistic segmentation loss *ℒ*_*seg*_ can be formulated as Equation 2, where *Ŷ*_*DEM*_ is the predicted DEM segmentation and vol(*Ŷ*_*DEM*_,*Y*_*DEM*_) is the newly proposed volume loss to avoid over- and under-segmentation in relation to the future volume of WMH (discussed in the next section).

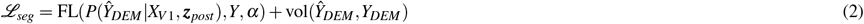

### Volume loss

To avoid over- and under-segmentation in relation to the future volume of WMH, a volume-loss (that is formulated as Equation 3) is added to Equation 2 as regularization term. The term keeps the predicted future volume of WMH (i.e., calculated from the predicted DEM (*Ŷ*_*DEM*_)) close to the reference future volume of the WMH (i.e., calculated from the ground truth DEM (*Y*_*DEM*_)) by using mean squared error (MSE). Note that only class *c* = 2 (for growing WMH) and class *c* = 3 (for stable WMH) that matter in the calculation of future volume of WMH at V2. A denominator of 1000 was used to estimate the volume of WMH in *ml* (i.e., as voxel dimensions are in *mm*^3^). In preliminary experiments, we found that having the same weights for segmentation loss of FL(*P*(*Ŷ*_*DEM*_|*X*_*V*1_, ***z***_*post*_),*Y, α*) and volume loss of vol(*Ŷ*_*DEM*_,*Y*_*DEM*_) in Equation 3 produced the best results for predicting both future volume and spatial dynamic changes of WMH.

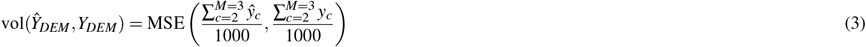

### Probabilistic loss

We used *Kullback-Leibler* Divergence score (*𝒟*_*KL*_) in the training process for training the Prior Net and Posterior Net. In this setting, Prior Net and Posterior Net were trained together with the Segmentation Net for predicting the DEM. Let *Q* be the posterior distribution from the Posterior Net and *P* be the prior distribution from the Prior Net. The difference between the posterior distribution *Q* and the prior distribution *P* is described by *𝒟*_*KL*_ in Equation 4 where *X*_*V*2_ is the T2-FLAIR at V2, *Y*_*DEM*_ is the true DEM, and *X*_*V*1_ is the T2-FLAIR at V1. Based on our preliminary experiments, the dimension for both ***z***_*post*_ and ***z***_*prior*_ is 4 (smaller than the original paper^26^ which used 6).

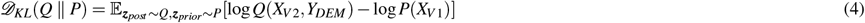

### Adversarial loss

Similar to a previous study^25^, the original adversarial loss proposed by^39^ was slightly modified by adding a segmentation loss (*ℒ*_*seg*_) so that the Segmentation Net was also optimised to produce better segmentation result. Similar to the original paper^39^, the Segmentation Net aims at minimising Equation 5 while the discriminator network aims at maximising it. In Equation 5, *G* is the Segmentation Net, *D* is the discriminator network, *y∼* (*X*_*V*1_, *X*_*V*2_,*Y*_*DEM*_) is the joint distribution of T2-FLAIR MRI at V1 and V2 and ground truth DEM (i.e., *X*_*V*1_, *X*_*V*2_, and *Y*_*DEM*_ respectively), *x∼* (*X*_*V*1_, *X*_*V*2_,*Ŷ*_*DEM*_) is the joint distribution of T2-FLAIR MRI at V1 and V2 and predicted DEM (i.e., *X*_*V*1_, *X*_*V*2_, and *Ŷ*_*DEM*_ respectively), 𝔼_*y*_ *∼Y_GAN_* is the expected value over *Y*_*GAN*_, and 𝔼_*x*_ is the expected value over *X*_*GAN*_.

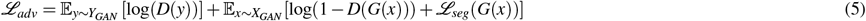

### Evaluation measurements

In this study, we used the following evaluation measurements to assess the performance of all configurations.

1. **Volume error** measures how close the predicted WMH volumes are with the real WMH volumes at the follow-up assessment (V2). This is the main performance measurement. Volume error can be calculated by using Equation 6 where 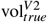 is the true volume of WMH V2, 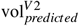 is the predicted volume of WMH at V2, and 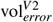 is the volume error.

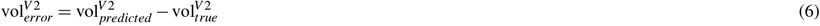
2. **Accuracy of prediction** assesses how good our proposed models predict WMH evolution for all patients (i.e., growing or shrinking). Accuracy of prediction for growing and shrinking WMH (i.e., subjects with growing and shrinking WMH are correctly predicted to have growing and shrinking WMH respectively) is calculated by the Equations 7 and 8 respectively. *N*_GRW_ and *N*_SHR_ are the number of subjects in our dataset who have growing and shrinking WMH. Whereas, *P*_GRW_ and *P*_SHR_ are the number of subjects correctly predicted as having growing and shrinking WMH.

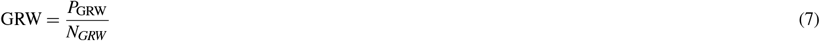

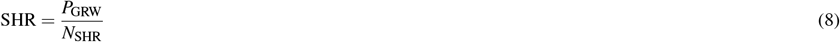
3. **Estimated volume interval (EVI)** measures the deviation of the predicted WMH volume at follow-up (V2) from the lowest and highest possible predicted volumes of WMH^25^. The lowest and highest possible predicted volumes of WMH at V2 are estimated by ignoring the prediction channel for growing WMH and shrinking WMH respectively. In other words, the lowest possible volume of WMH (dubbed as Minimum Volume Estimation or ‘MinVE’) is assumed to occur when there are no growing WMH in the patient’s brain. Whereas, the highest possible volume of WMH (dubbed as Maximum Volume Estimation or ‘MaxVE’) is assumed to occur when there are no shrinking WMH in the patient’s brain. There are 3 metrics in this evaluation: “CP” which stands for “Correct Prediction” (calculated by using Equation 9), “CPinEVI” which stands for “Correct Prediction in Estimated Volume Interval” (calculated by using Equation 10), and “(CP+WP)inEVI” which stands for “Correct Prediction + Wrong Prediction but still in EVI” (calculated by using Equation 11). In these equations, 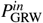 and 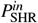 are the number of subjects that are correctly predicted as having growing and shrinking WMH and have their estimated volumes of WMH at V2 are located between ‘MinVE’ and ‘MaxVE’. Whereas, *P*^*in*^ is the number of subjects whose estimated volumes of WMH at V2 are located between ‘MinVE’ and ‘MaxVE’.

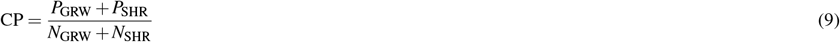

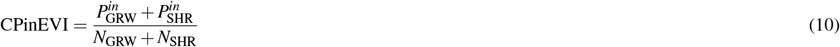

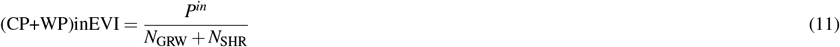
4. **Spearman correlation with Prins clinical scores** The clinical scoring system for progression of WMH, known as Prins visual scores^38^, gives a +1 for each WMH cluster that increases or appears *de nuovo* in a subsequent scan compared with a previous scan in the periventricular or deep white matter of each lobe (i.e., frontal, parietal, temporal and occipital), -1 if a reduction in volume or disappearance of a WMH cluster is detected, and 0 if no change can be appreciated. For our evaluation, we summed the overall scores in each region to obtain a total Prins score. We calculate the Spearman correlation between the total Prins scores and the spatial volume growth, shrinkage, and overall change that each scheme outputs.
5. **Spatial agreement** between predicted and ground truth DEM is measured by the Dice similarity coefficient (DSC)^49^. Higher values of DSC mean better performance. DSC can be calculated by using Equation 12, where *TP* is true positive, *FP* is false positive and *FN* is false negative. This is a secondary performance measurement as predicted future WMH volumes at V2 are calculated from segmentation masks.

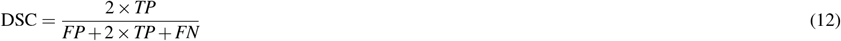
6. **Uncertainty quantification and correlation analysis** to measure correlation between uncertainty values in predicted DEM and DSC values, is calculated as the Cross Entropy (CE) between the mean sample and all samples as per Equation 13 where *γ* is the uncertainty map, *s* is a set of predictions from an input, *ŝ* is the mean sample of set *s*, CE is the cross entropy function, and 𝔼 is the expected value function.

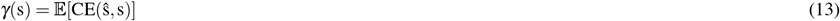

## Results and Discussion

In this section, we show and discuss the results of the evaluations using the four performance measurements, namely volume based evaluation, qualitative/visual evaluation, spatial agreement evaluation,uncertainty quantification, and correlation with clinical visual scores.

### Evaluation against clinical visual scores of WMH progression

Figure 4 shows the results from calculating the non-parametric correlations between Prins clinical visual scores and the spatial volume growth and shrinkage from each scheme. The spatial growth from all models correlated with Prins scores, with the output from PUNet-vol showing the highest correlation following the ground truth (Spearman rho = 0.40 and 0.58 respectively). This correlation slightly improved (i.e., to Spearman rho = 0.42) when attention was incorporated in the scheme. Prins showed net shrinkage for only six patients, as shrinkage in individual clusters were nullified by growth in others. The ground truth showed the worst correlation with Prins in terms of shrinkage (Spearman rho = 0.45), followed by PUnet-wSL (Spearman rho = 0.47 without attention and 0.50 with it). The highest correlation values in shrinkage were achieved with PUNet-vol without attention (Spearman rho = 0.59), and PUNet with attention (Spearman rho = 0.60). In general, in models without attention WMH shrinkage and growth correlated better with Prins than when attention was used. In line with a previous study^50^, the spatial net change did not correlate with Prins, neither improving when attention was used.

### Results on predicting future volumes of WMH

WMH volume change is an important clinical feature for clinical research and could be an important biomarker if available for clinical practice. Hence, we evaluated how well WMH volume at V2 (1 year later) can be estimated by using our proposed models. Table 1 shows the prediction accuracy of WMH volumetric progression (i.e., whether WMH volume will grow or shrink at V2 for each patient) calculated using Equations 7 and 8, the estimated volume interval (EVI) calculated using Equations 9, 10, and 11, and the volumetric error calculated using Equation 6.

**Table 1.**
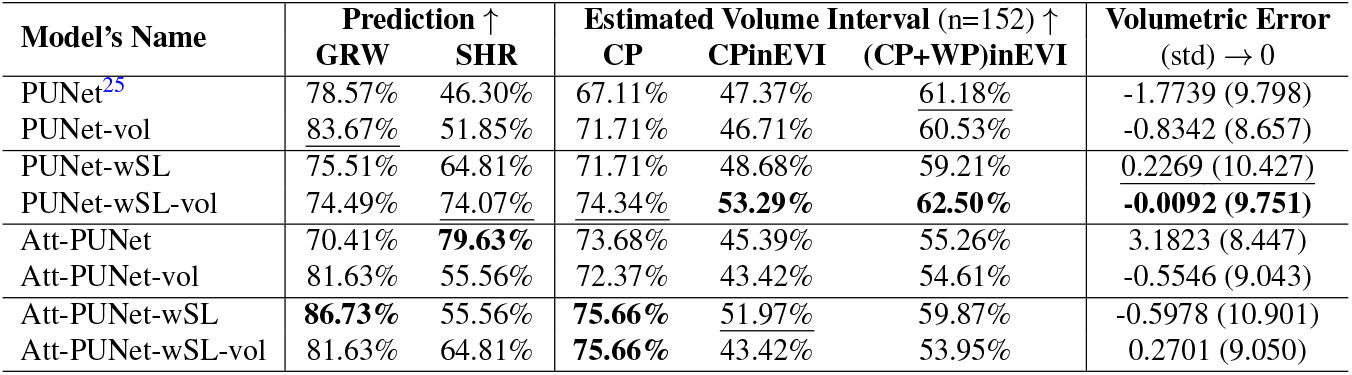
Volume based evaluation for all models evaluated. There are 98 patients with growing (GRW) and 54 with shrinking (SHR) volume of WMH. “CP” stands for “Correct Prediction”, “CPinEVI” stands for “Correct Prediction in Estimated Volume Interval”, and “(CP+WP)inEVI” stands for “Correct Prediction + Wrong Prediction but still in EVI”. Symbol indicates that higher values are better while symbol → 0 indicates that values closer to 0 are better. The best and second best values for each evaluation measurements are written in bold and underlined respectively.

| Model’s Name | Prediction ↑ | | Estimated Volume Interval (n=152) ↑ | | | Volumetric Error<br>(std) $\rightarrow 0$ |
| --- | --- | --- | --- | --- | --- | --- |
|  | GRW | SHR | CP | CPinEVI | (CP+WP)inEVI |  |
| PUNet<sup>25</sup> | 78.57% | 46.30% | 67.11% | 47.37% | 61.18% | -1.7739 (9.798) |
| PUNet-vol | 83.67% | 51.85% | 71.71% | 46.71% | 60.53% | -0.8342 (8.657) |
| PUNet-wSL | 75.51% | 64.81% | 71.71% | 48.68% | 59.21% | 0.2269 (10.427) |
| PUNet-wSL-vol | 74.49% | 74.07% | 74.34% | **53.29%** | **62.50%** | **-0.0092 (9.751)** |
| Att-PUNet | 70.41% | **79.63%** | 73.68% | 45.39% | 55.26% | 3.1823 (8.447) |
| Att-PUNet-vol | 81.63% | 55.56% | 72.37% | 43.42% | 54.61% | -0.5546 (9.043) |
| Att-PUNet-wSL | **86.73%** | 55.56% | **75.66%** | 51.97% | 59.87% | -0.5978 (10.901) |
| Att-PUNet-wSL-vol | 81.63% | 64.81% | **75.66%** | 43.42% | 53.95% | 0.2701 (9.050) |

As Table 1 shows, PUNet-wSL-vol performed better than the rest of the models producing either the best or second best results for almost all evaluation metrics except predicting growing WMH (GRW). There were more patients with net growing WMH than with net shrinking WMH in the dataset, thus hinting to a possible bias by the other models towards growing WMH. Reduction in WMH volume was mainly observed in patients with high WMH volume (see Figure 3B).

**Figure 3.**
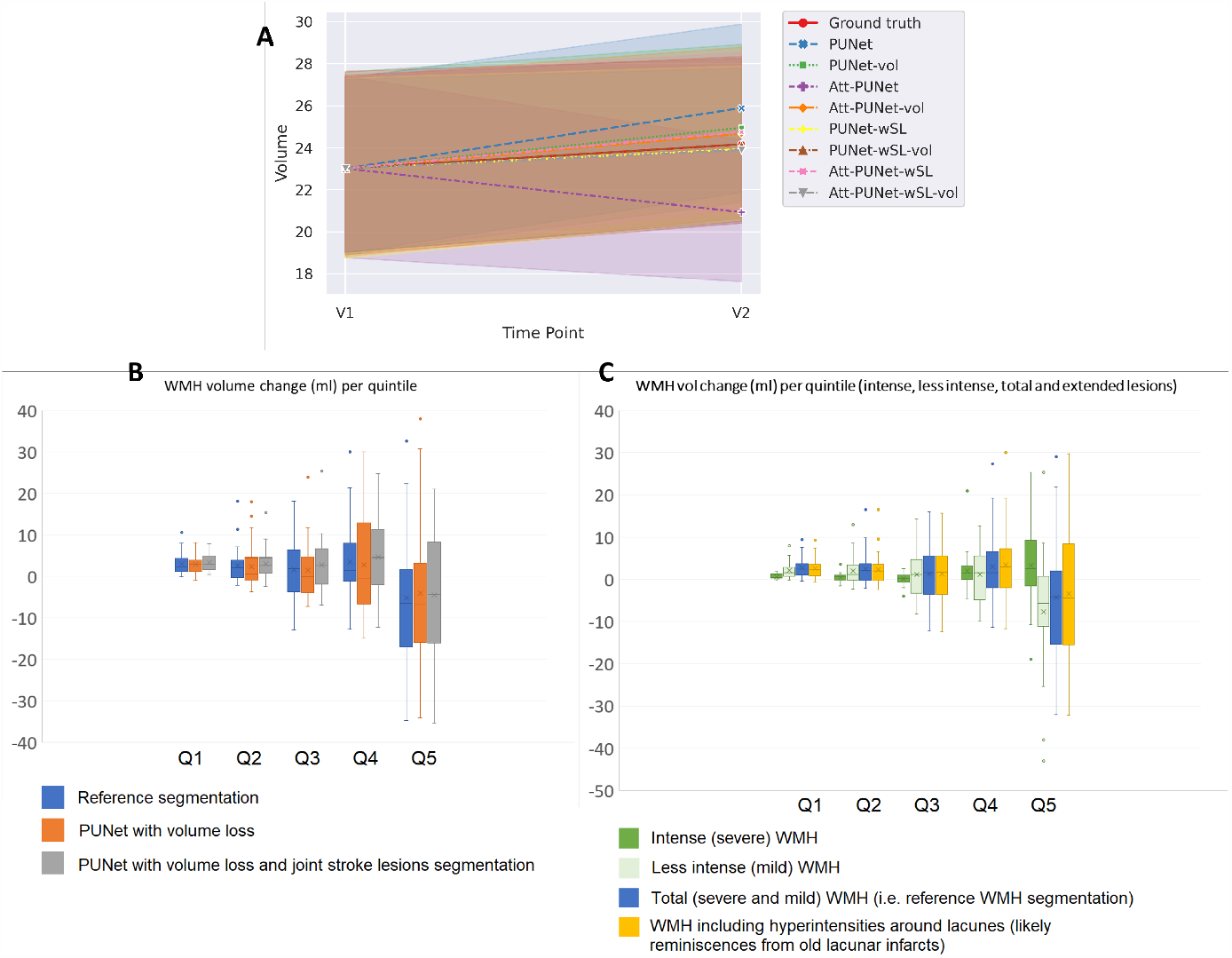
**A:** Average progression of WMH volume (*ml*) from V1 to V2 (1 year) for Ground truth and all tested models/configurations. **B:** Volumetric WMH change in *ml* (vertical axes) for patients grouped by quintiles (horizontal axes) depending on their WMH volume at baseline.

**Figure 4.**
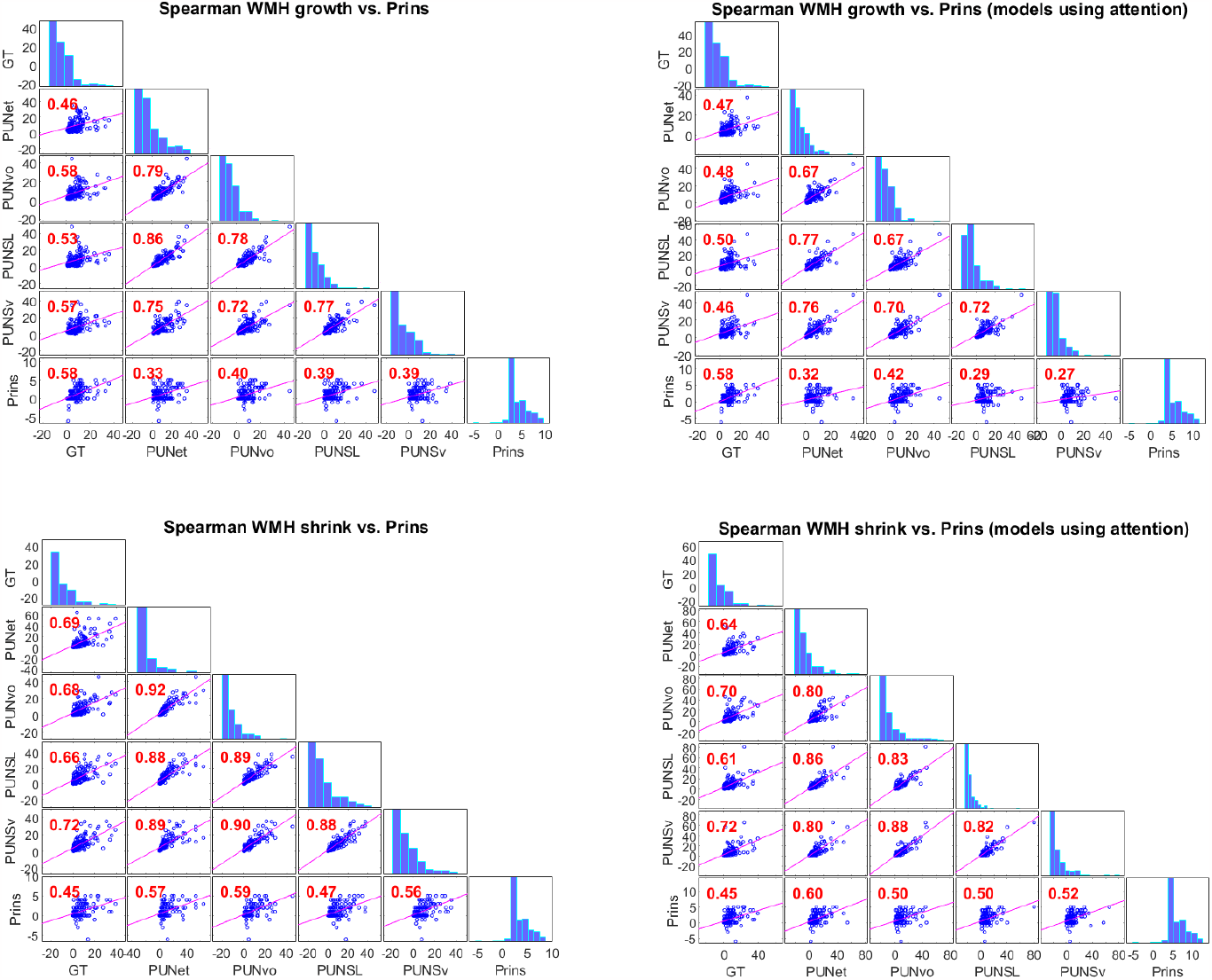
Spearman correlations between the spatial growth (above) and shrinkage (below) from each scheme and Prins overall scores. GT: ground truth, PUNet: Probabilistic UNet, PUNvo: Probabilistic UNet adding a volume loss to the cost function (PUNet-vol), PUNSL: Probabilistic UNet that outputs stroke lesion (SL) and WMH masks, PUNSv Probabilistic UNet that outputs SL and WMH and uses volume loss added to the cost function, Prins: total (summed) clinical scores

As Figure 3A shows, the average progression of WMH volume from V1 to V2 (in *ml*) was well estimated by PUNet-wSL-vol (i.e., brown dashed line representing PUNet-wSL-vol is coincident with the red line representing the ground truth). In general, as expected, models trained using volume loss (Equation 3) (i.e., PUNet-vol (green line), Att-PUNet-vol (orange line), PUNet-wSL-vol (brown line), and Att-PUNet-wSL-vol (grey line)) produced more accurate progression of WMH volume from V1 to V2 than those which did not use volume loss during training. Of note, however, PUNet-wSL (yellow line) and Att-PUNet-wSL (pink line), had lines close to the red line of the ground truth. Overall, models jointly segmenting stroke lesions and WMH improved the estimation of future volume of WMH at V2.

To further analyse the accuracy of the winner scheme in estimating the WMH volume change, we grouped the patients in quintiles according to their WMH volume at baseline and calculated the WMH change produced by the reference (i.e., ground truth) segmentation, and the PUNet model using volume loss with and without jointly segmenting the DEM and the stroke lesions. As can be appreciated from Figure 3B, the scheme that jointly segmented the stroke lesions and the DEM of WMH change produced mean, median and a distribution of WMH volume change values across the sample more similar to those from the reference segmentation, than the scheme that only segmented the DEM of WMH change for all but the highest quintile.

We also divided the reference WMH segmentations into intense and less intense WMH as per^44^, and considered an ‘extended’ WMH volume that included the WMH surrounding lacunes, thought to be reminiscences of old small subcortical infarcts (see Figure 3C). It can be observed that the volume output from the scheme that jointly segmented the stroke lesions with the DEM of WMH change resulted strikingly similar to the one produced by this ‘extended’ WMH segmentation (see gray and yellow box plots in Figures 3B and 3C, respectively), especially for patients in the highest quintile. Patients in this quintile exhibit a high burden of WMH surrounding lacunes and coalescing with previous strokes. Therefore, it is expected that not only AI schemes, but also experts would consider all hyperintensities as part of the white matter disease in absence of any other sequence or clinical information from this patient group. It can be also seen that the reference WMH change (i.e., blue box plot in the same figure) is mainly determined by the less intense WMH change (i.e., pale green box plot), therefore explaining the difficulty in obtaining accurate growth and shrinking spatial estimates and putting into question the accuracy in the spatial estimates of the ground truth segmentations given the degree of observer-dependent manual input they had.

### Spatial agreement evaluation

Spatial agreement evaluation was performed to see whether closer predicted future volumes of WMH to the reference future WMH volumes is followed by higher spatial agreements between predicted DEM and ground truth DEM or not. Table 2 shows performances of all tested configurations using DSC (Equation 12). The best and second best measurement values for each DEM label are written in bold and underlined respectively. Note that the ‘Changing’ refers to shrinking and growing WMH combined together as one label, ‘Average’ is calculated by averaging DSC values of ‘Shrinking’, ‘Growing’, and ‘Stable’, and ‘Stroke Lesions’ is only available when joint segmentation of WMH DEM and stroke lesions are performed.

**Table 2.** Dice similarity coefficient (DSC) for all model configurations. Symbol ↑ indicates that higher values are better. The best and second best measurement values for each category of WMH are written in bold and underlined respectively.

| Model's Name | Dice Similarity Coefficient (DSC) ↑ | | | | | |
| --- | --- | --- | --- | --- | --- | --- |
|  | Shrinking | Growing | Stable | Average | Changing | Stroke Lesions |
| PUNet<sup>25</sup> | 0.2132 | <u>0.2137</u> | 0.6385 | 0.3551 | 0.3633 | - |
| PUNet-vol | 0.2107 | **<u>0.2232</u>** | **<u>0.6439</u>** | 0.3593 | 0.3642 | - |
| PUNet-wSL | <u>0.2217</u> | 0.2130 | <u>0.6437</u> | <u>0.3595</u> | **<u>0.3719</u>** | 0.4499 |
| PUNet-wSL-vol | **<u>0.2290</u>** | 0.2112 | 0.6392 | **<u>0.3598</u>** | <u>0.3681</u> | 0.4281 |
| Att-PUNet | 0.2211 | 0.1796 | 0.6302 | 0.3437 | 0.3510 | - |
| Att-PUNet-vol | 0.2078 | 0.1981 | 0.6315 | 0.3458 | 0.3471 | - |
| Att-PUNet-wSL | 0.1968 | 0.2045 | 0.6240 | 0.3417 | 0.3543 | <u>0.5338</u> |
| Att-PUNet-wSL-vol | 0.1960 | 0.2077 | 0.6322 | 0.3453 | 0.3536 | **<u>0.5430</u>** |

From Table 2, we can see that joint segmentation of DEM and stroke lesions with volume loss (PUNet-wSL-vol) produced the best segmentation results based on DSC for ‘Shrinking’ (0.2290) and ‘Average’ (0.3598). Furthermore, we can see that joint segmentation of DEM and stroke lesions by PUNet-wSL (i.e., without volume loss) and PUNet-wSL-vol (i.e., with volume loss) produced either the best or second best DSC values for all categories of DEM, except for ‘Growing’ and ‘Stable’ WMH (achieved by the original models of PUNet and PUNet-vol). On the other hand, models with auxiliary input of probabilistic maps of WMH change, (i.e., Att-PUNet, Att-PUNet-vol, Att-PUNet-wSL, and Att-PUNet-wSL-vol) failed to improve the performance of the DEM segmentation while improved the performance of ‘Stroke Lesions’ segmentation. Furthermore, models trained using volume loss (i.e., PUNet-vol, Att-PUNet-vol, PUNet-wSL-vol, and Att-PUNet-wSL-vol) produced better DSC values on ‘Average’, which indicates that the volume loss impacted positively in the task of estimating the DEM of WMH.

DSC is influenced by TP, FP, and FN counts between ground truth mask and predicted segmentation, but TP, FP, and FN counts are highly imbalance in the segmentation of brain lesions. To provide a better illustration of the relationship between DSC and corresponding TP, FP, and FN counts, we present the confusion matrices and a table compiling these values from the ‘Shrinking’ WMH and ‘Growing’ WMH labels obtained from PUNet-vol and PUNet-wSL-vol configurations (Figure 5 and Table 3 respectively). Figure 5 contains the number of segmented voxels corresponding to each label (*n*) from all patients in the testing set, false negative rate (*fnr*), false positive rate (*fpr*), true positive rate (TPR), and positive predictive value (PPV). Table 3 compiles values of DSC, PRE, REC, FN, and FP for the ‘Shrinking’ WMH and ‘Growing’ WMH labels from both PUNet-vol and PUNet-wSL-vol configurations. From both, Figure 5 and Table 3, we can see that PUNet-vol produced higher PRE value for ‘Shrinking’ WMH with lower FP counts than PUNet-wSL-vol. But PUNet-vol produced lower PRE value for ‘Growing’ WMH as it produced higher FP counts than PUNet-wSL-vol in this label/category.

**Table 3.**
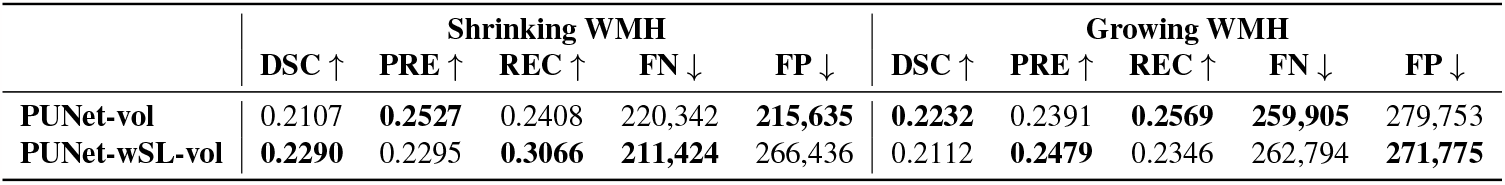
Comparison of DSC, PRE, and REC values to FN and FP counts for PUNet-vol and PUNet-wSL-vol configurations. Symbols *↑* and *↓* indicate that higher and lower values are better respectively.

|  | Shrinking WMH |  |  |  |  | Growing WMH |  |  |  |  |
| --- | --- | --- | --- | --- | --- | --- | --- | --- | --- | --- |
| | DSC ↑ | PRE ↑ | REC ↑ | FN ↓ | FP ↓ | DSC ↑ | PRE ↑ | REC ↑ | FN ↓ | FP ↓ |
| PUNet-vol | 0.2107 | **<u>0.2527</u>** | 0.2408 | 220,342 | **<u>215,635</u>** | **<u>0.2232</u>** | 0.2391 | **<u>0.2569</u>** | **<u>259,905</u>** | 279,753 |
| PUNet-wSL-vol | **<u>0.2290</u>** | 0.2295 | **<u>0.3066</u>** | **<u>211,424</u>** | 266,436 | 0.2112 | **<u>0.2479</u>** | 0.2346 | 262,794 | **<u>271,775</u>** |

**Figure 5.**
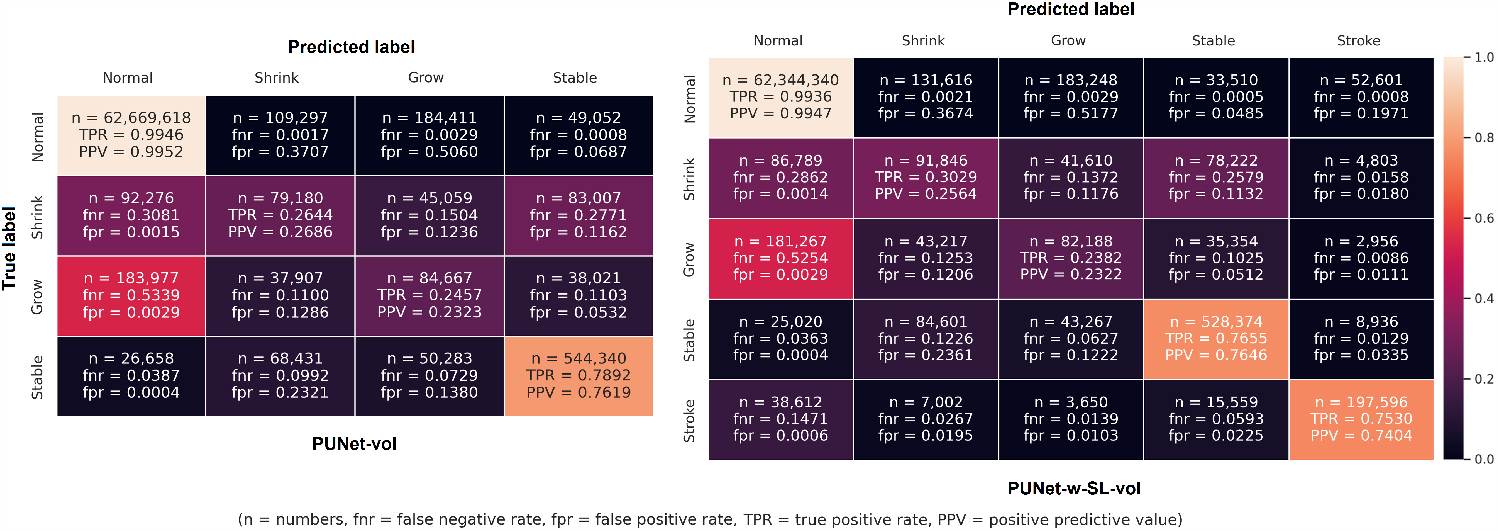
Confusion matrices for all labels produced by PUNet-vol and PUNet-wSL-vol configurations from all subjects. Abbreviation *n* stands for number of segmented voxels which can be used to calculate false negative rate (*fnr*), false positive rate (*fpr*), true positive rate (TPR), and positive predictive value (PPV). Note that TPR and *fnr* are calculated horizontally for each row (true label of DEM). On the other hand, PPV and *fpr* are calculated vertically for each column (predicted label of DEM).

Confusion matrices in Figure 5, show a high level of uncertainty between ‘Growing’ WMH and ‘Normal’ brain tissues as more than 50% of the ‘Growing’ WMH identified in the ground truth DEM were wrongly predicted as ‘Normal’ tissues (i.e., under-segmentation of ‘Growing’ WMH which leads to higher *fnr* in the confusion matrix) by PUNet-vol and PUNet-wSL-vol configurations with *fnr* = 0.5339 and *fnr* = 0.5254 respectively. In extended experiments, all proposed configurations were observed producing the same level of under-segmentation for ‘Growing’ WMH. In general, areas of ‘Growing’ WMH are difficult to be differentiated from ‘Normal’ brain tissues due to the high level of uncertainty between these two classes. Overall, for the model that jointly segmented the stroke lesions and the WMH, mean DSC values were slightly better in this sample.

Although the combined segmentation of WMH and stroke lesions is not the main focus of this study, it must be noted that the state-of-the-art joint segmentation method for WMH and stroke lesions (i.e., sub-acute and chronic as per in the present dataset)^51^, which used a UResNet configuration, reported a mean (SD) Dice equal to 0.4 (0.252) for stroke lesions segmentation, lower than any of our joint-segmentation schemes (see Table 2).

### Qualitative/visual evaluation of spatial agreement between ground truth and predicted DEMs

Figures 6A and 6B show examples of the predicted DEM segmentation from PUNet-wSL-vol and PUNet-vol and their corresponding DEM ground truth forpatients with high and low DSC values on ‘Average’ respectively. PUNet-wSL-vol and PUNet-vol were chosen for qualitative/visual evaluation as they produced the best and second best DSC values on ‘Average’ (See Table 2). Figure 6A shows that PUNet-wSL-vol, which jointly segments WMH DEM and stroke lesions, produced better segmentation results than PUNet-vol, which exhibits a high level of uncertainty in predicting shrinking and growing WMH. Confusion matrices in Figure 5 show that PUNet-wSL-vol lowered this uncertainty by producing lower rates of *fnr* (and their corresponding FN counts (*n*)) for shrinking and growing WMH) in most cases. Figure 6B illustrates cases where low DSC values of predicted WMH DEM were caused mostly by two reasons: low WMH volume at V1 (patient and MSSB172) and brain MRI artefacts (patient MSSB211). Based on our observations, these two problems were relevant throughout the sample in our evaluations.

**Figure 6.**
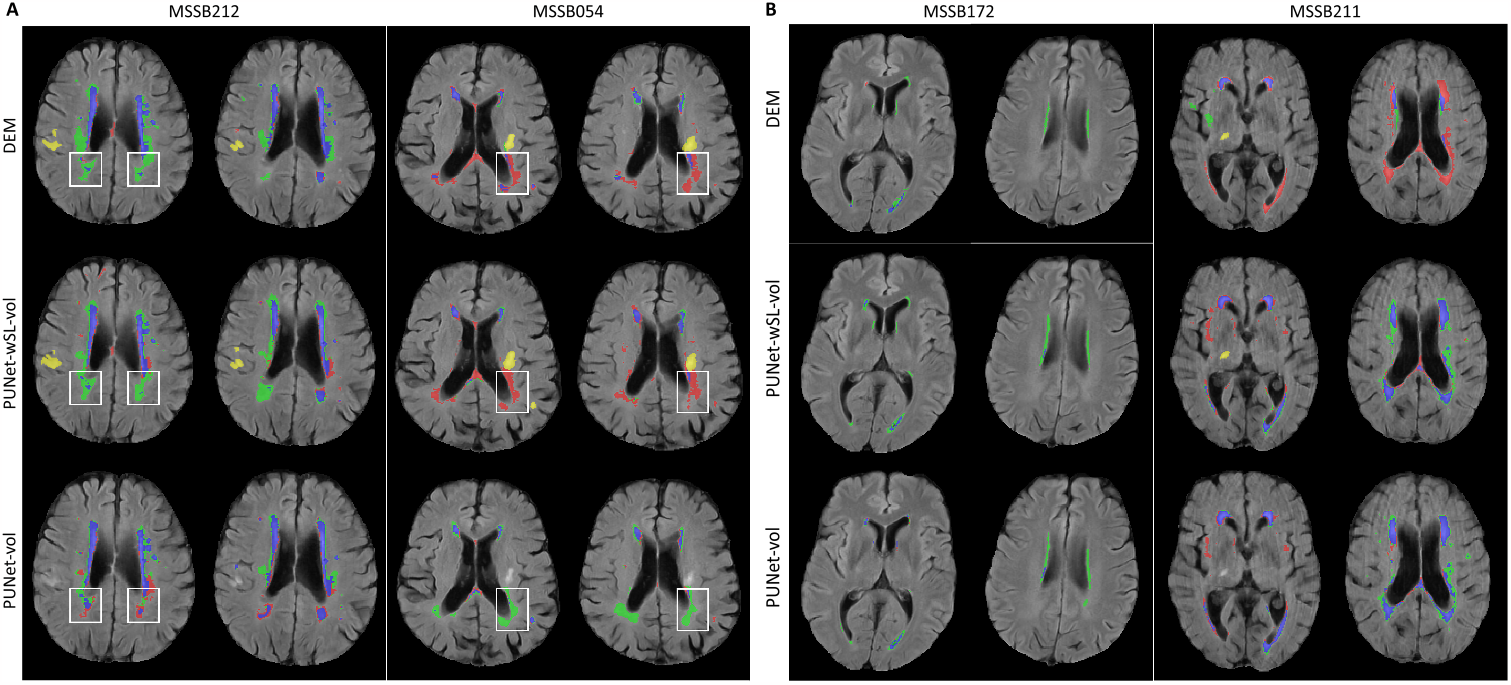
**A:** Examples of predicted DEM produced by PUNet-wSL-vol and PUNet-vol and their corresponding DEM ground truth from subjects with high DSC values on average. **B:** Examples of predicted DEM produced by PUNet-wSL-vol and PUNet-vol and their corresponding DEM ground truth from subjects with low DSC values on average. **A and B:** Red represents shrinking WMH, green represents growing WMH, blue represents stable WMH, and yellow represents stroke lesions. Obvious improvements are highlighted in white rectangles.

### Uncertainty quantification

As all configurations evaluated are based on the Probabilistic U-Net, uncertainty for each label in the DEM was quantified by predicting DEM for each subject multiple times. In this study 30 different DEM predictions were generated from 30 samples of *z*_*prior*_ from Prior Net for each input data/patient. From these 30 DEM predictions per patient data, uncertainty was calculated as the Cross Entropy (CE) between probability values from all DEM predictions and its average as written in Equation 13.

Figure 7 shows the uncertainty maps for all DEM labels produced by the model that generated the best DSC ‘Average’ value, PUNet-wSL-vol, for the whole brain and inside the predicted DEM for a patient. From the uncertainty maps for the whole brain, we can see that the uncertainties for shrinking and growing WMH encompass larger brain areas than for stable WMH. This finding supports results from evaluating the spatial agreement between ground truth and the models’ outputs, indicating lower accuracy in the predictions of changing WMH (i.e., shrinking and growing) compared to the predictions of stable WMH. The example shown in Figure 7A has incorrect areas showing uncertainty in the ‘Shrinking’ label (e.g., in the frontal cortex and in the septum), owed mainly to hyperintense flow artefacts.

**Figure 7.**
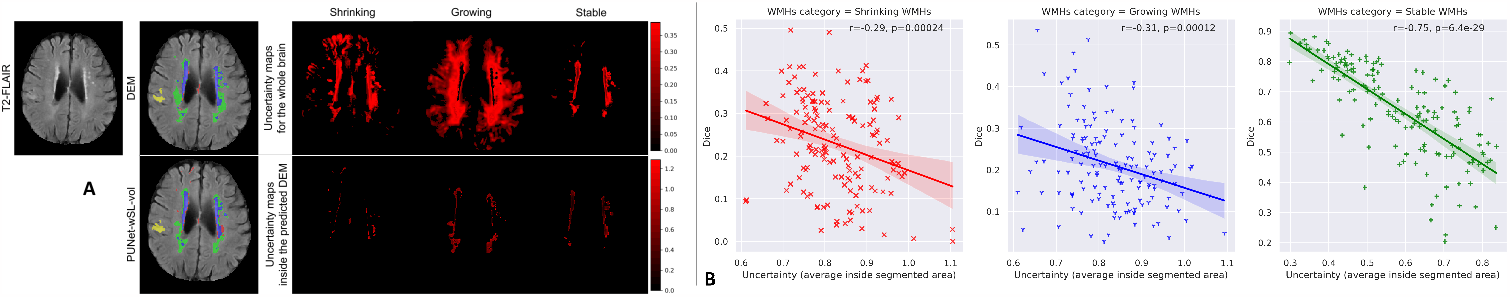
**A:** Uncertainty maps produced by PUNet-wSL-vol from subject MSSB212. **B:** Correlation between the average of uncertainty values inside the predicted DEM and DSC values of predicted DEM produced by PUNet-wSL-vol.

Interestingly, in the uncertainty maps, the uncertainty values inside DEM labels of shrinking and growing WMH are higher than those inside stable WMH, a consistent finding from this evaluation. This is in-line with a previous analysis^50^ that showed WMH progression and disappearance being associated with the areas of ill-defined subtle or “less intense” WMH, largely identified as indicative of pre-(and post-) lesional changes. As expected, and Figure 7B shows, the uncertainty values inside the predicted DEMs and the DSC values produced by PUNet-wSL-vol are negatively correlated for each DEM label (i.e., shrinking, growing and stable WMH). However, only for stable WMH (r=0.75) higher DSC values of DEM labels can be predicted by having lower uncertainty values inside the predicted DEM and vice versa. Plots of the correspondence in shrinking and growing labels show a wide spread in DSC values especially among those with uncertainty values between 0.7 and 0.98, that would make any inference of the predictive power of one magnitude over the other inaccurate.

### Conclusion

This study proposed deep learning models based on the Probabilistic U-Net^26^ architecture trained incorporating stroke lesions information into the models and using volume loss as additional loss for improving the quality of predicted future volume of WMH and disease evolution map (DEM) of WMH. Probabilistic U-Net was chosen as the baseline method because a preliminary study showed that it performed better than the U-Net^25^.

We proposed three different approaches for incorporating stroke lesions information into Probabilistic U-Net models. These are **(1)** joint segmentation of DEM and stroke lesions, **(2)** use of probabilistic maps of WMH change in relation to stroke lesions’ locations, and **(3)** combination of **(1)** and **(2)**. We proposed to incorporate stroke lesions information into deep learning models to predict WMH evolution because stroke is commonly associated with the evolution of WMH^3^. Based on the results from the various experiments, joint segmentation of DEM and stroke lesions (approach **(1)**) was the most effective approach to improve the quality of predicted DEM of WMH in all evaluations, while being also simpler and more straightforward than the other approaches evaluated in this study. The introduction of a volume loss as an additional loss to the scheme substantially improved the quality in predicting the DEM of WMH in terms of volume, correlation with clinical scores of WMH progression, and in terms of spatial agreement.

This study shows that 1) incorporating factors that have been commonly associated with WMH progression (i.e., stroke lesions information) is crucial to produce better prediction of DEM for WMH from brain MRI; 2) the best method for incorporating associated factors that can be extracted from the same data/image modality involves performing multi-task learning; and 3) in patients with vascular pathology, a multi-class segmentation of brain features resulting from symptomatic (i.e. stroke) and asymptomatic (i.e., WMH) vascular events generates better results consistent with clinical research. In this study, as stroke lesions appear on the same T2-FLAIR MRI sequence as WMH, we performed joint segmentation of DEM for WMH and stroke lesions. However, previous clinical studies have shown that there are other non-image risk factors and brain features that have been commonly associated with the progression and evolution of WMH, like age^8^, ventricular enlargement^52,53^, and brain atrophy^54^. Thus, more (image and non-image) factors could be incorporated in future studies to further improve the quality of predicted DEM of WMH, although the best way to incorporate non-image factors to the prediction model remains to be found.

This study also has limitations to overcome in future works. The dataset was small in size impeding a quantitative in-depth analysis of the models’ performance in different patient subgroups, e.g., patients stratified by age and sex, patients grouped by stroke subtype, etc. Thus, subgroup analyses were carried out visually and volumetrically, not spatially. By using only data from patients presenting to a clinic with a mild-to-moderate stroke, the generalisability of the proposed approach can be questioned. Therefore, further evaluation in a wider and more heterogeneous sample will be needed. The use of DSC in the evaluation needed the binarisation of the probabilistic outputs from the models. Limitations in the use of DSC have been recently acknowledged^55^. However, it must be noted that ground truth segmentations are also binary, and observer-dependent. By using different quality control metrics in a comprehensive analysis we have overcome the limitations that the analysis of the spatial agreement using DSC poses. A probabilistic metric allowing spatial analyses of segmentation results is needed. Also, we used probabilistic maps of WMH change for strokes in the lentiform nucleus and centrum semiovale based on findings from a clinical study. However, the same clinical study specified that it was not possible to ascertain WMH evolution and distribution for patients with the stroke in other regions like thalami and midbrain or brain stem due to the limited sample of patients with infarcts in those regions. Incorporating findings for more powered studies would be necessary to conclude about the usefulness of incorporating attention maps to the AI schemes. Finally, various schemes for estimating uncertainty in segmentation/classification tasks have recently emerged^56,57^, which would be worth exploring in the future for estimating WMH evolution.

## Acknowledgements

Funds from JSPS (Kakenhi Grant-in-Aid for Research Activity Start-up, Project No. 20K23356) (MFR); RIKEN’s Special Postdoctoral Researchers (SPDR) program (MFR); Row Fogo Charitable Trust (Grant No. BRO-D.FID3668413) (MCVH); Wellcome Trust (patient recruitment, scanning, primary study Ref No. WT088134/Z/09/A); Fondation Leducq (Perivascular Spaces Transatlantic Network of Excellence); EU Horizon 2020 (SVDs@Target); and the MRC UK Dementia Research Institute at the University of Edinburgh (Wardlaw programme) are gratefully acknowledged. This research was also supported by the program for Brain Mapping by Integrated Neurotechnologies for Disease Studies (Brain/MINDS) from the Japan Agency for Medical Research and Development AMED (JP15dm0207001). Library access provided by the Faculty of Computer Science, Universitas Indonesia is also gratefully acknowledged.

## Author contributions statement

M.F.R and M.V.H. conducted experiments (including conceptualization, methodology, software, experiments, and evaluations).

S.M. and J.W. performed data curation and clinical analysis. H.S. conducted methodology development and analysis.

## Additional information

**Accession codes:** https://github.com/febrianrachmadi/probunet-gan-vie; **Competing interests:** No conflict of interests.

The corresponding author is responsible for submitting a competing interests statement on behalf of all authors of the paper.

This statement must be included in the submitted article file.

## Footnotes

1 https://datashare.ed.ac.uk/handle/10283/3934

